# Sterilization and material compatibility of 3D-printed devices for cell culture

**DOI:** 10.64898/2026.08.03.742522

**Authors:** Matthew J. Footer, Nathan M. Belliveau

## Abstract

Three-dimensional (3D) printing is increasingly used in cell biology to prototype custom cell culture devices, microscopy chambers, and other experimental hardware, but practical guidance on sterilization and material compatibility remains limited. Here, we develop and validate a low-temperature paraformaldehyde vapor sterilization method for fused filament fabrication (FFF) components that avoids heat-induced deformation. Using bacterial challenge assays with *Escherichia coli* and *Geobacillus stearothermophilus*, we confirm effective sterilization of 3D printed components. We also assess the effects of common filaments and adhesives on HL-60 human cell growth to identify materials suitable for cell culture workflows. Most untreated plastics were well tolerated over 48 hours, whereas some formaldehyde-treated materials required post-treatment with ammonia vapor to restore compatibility. Together, these results provide a practical framework for sterilizing and deploying 3D-printed materials in cell culture and biological research.

**Multidisciplinary Abstract:** 3D printers allow laboratories to quickly make custom experimental equipment, but many printed materials cannot be sterilized using standard methods. We developed a low-temperature sterilization procedure using paraformaldehyde vapor and tested whether common printing materials and adhesives are compatible with cultured human cells. We identified several plastics, silicone-based adhesives and acrylic tapes that can be safely used after sterilization, while others require additional ammonia treatment. These results provide guidance for researchers using 3D printing to build custom laboratory and cell culture devices.

**Methods Summary:** A low-temperature paraformaldehyde vapor sterilization method for 3D-printed FFF components was developed and validated using bacterial challenge assays. Compatibility of common printing filaments, adhesives, and tapes with cell culture workflows was assessed by exposing HL-60 cells to untreated, sterilized, and ammonia- neutralized materials and quantifying cell growth by flow cytometry.

## Introduction

3D printing is now a well-established manufacturing method that has seen widespread adoption in teaching environments and research laboratories across academia and industry (1, 2). Even when a final device will ultimately be produced by injection molding or subtractive manufacturing (e.g., milling), 3D printing enables rapid iteration by allowing multiple prototypes to be built and tested before finalizing a design. This is especially valuable in research, where new tools and techniques are often integral to experimental progress. Online repositories of 3D models such as the NIH 3D Print Exchange (3) hosts thousands of printable models, ranging from protein structures and organisms to practical laboratory accessories.

In cell biology research, 3D printing is increasingly used to prototype and disseminate custom experimental hardware, including live-cell microscopy chambers, microfluidic platforms for cell handling and stimulation, and culture devices that interface with standard workflows or soft-lithography pipelines (4–9). In recent work (10, 11), we used 3D-printed assemblies that combined printed components with track-etch membranes and multiple adhesives that enabled us to screen millions of cells and isolate subsets based on motility phenotypes (Fig. 1). This has allowed us to extend the utility of genome-scale phenotypic screens to the study of cell motility, a process fundamental to diverse processes that include immune cell function, multicellular development, and the dissemination of cancer cells during metastasis.

**Figure 1.**
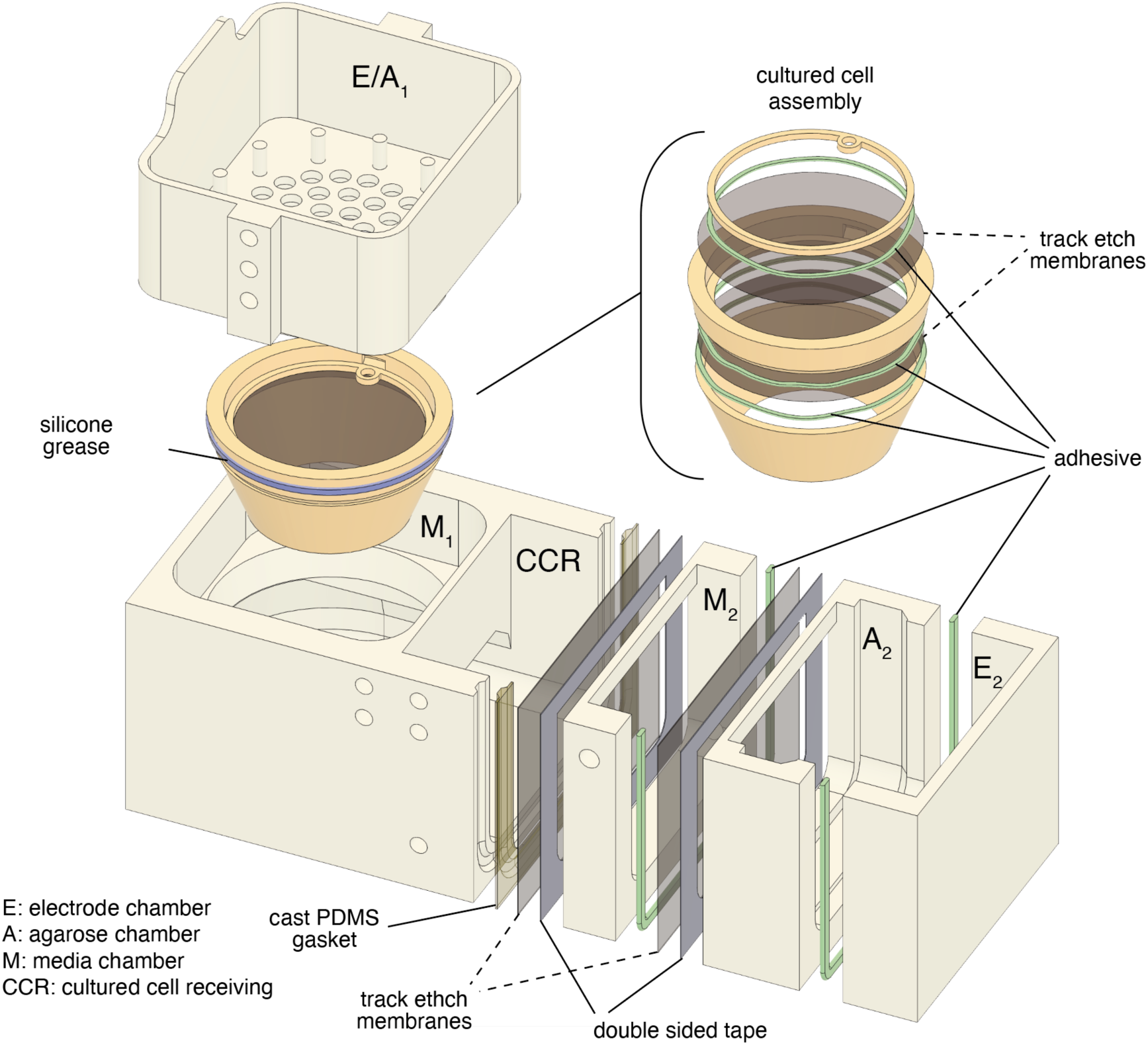
Example of an FFF-printed device used in cell-biological research. This exploded view shows a modular assembly used to isolate cultured cells according to their migratory response to an electric field. The white and cream components were printed in PLA. The device also incorporates silicone adhesive, double-sided tape, track-etched membranes, PDMS gaskets cast in 3D-printed molds, and silicone grease used as a temporary watertight seal. These materials were compatible with cell culture under the conditions tested. E, electrode chamber; A, agarose chamber; M, media chamber; CCR, cultured-cell receiving chamber. Electrodes, agarose, and culture medium are omitted for clarity.

One remaining challenge to the broader utility of these fabrication tools in biological research is that information about the compatibility of 3D printer plastics, adhesives, and other fabrication materials with biological systems remains limited. Consumer-grade 3D printer filament for fused filament fabrication (FFF) is often not a pure polymer, but a complex formulation containing additives such as lubricants, stabilizers, and biocides (12). Since printer filament is not typically produced with biological research applications in mind, printed devices may introduce unintended experimental perturbations if leachable chemicals or additives accumulate in biological solutions (13). Furthermore, many common filament materials cannot be readily sterilized using standard laboratory methods without heat-induced deformation, limiting their utility in longer-term cell culture experiments that require stringent contamination control.

In this work, we provide characterization of commonly available filament plastics and adhesives for use in cell biological research. We develop and validate a low-temperature paraformaldehyde vapor sterilization protocol compatible with 3D-printed parts. We then assess potential toxicity of fabricated components by measuring effects on proliferation of the human promyelocytic leukemia cell line HL-60, which is widely used in immune cell biology, drug screening, and toxicology (14–18). Together, these findings establish practical guidelines and compatible materials to help biological research groups incorporate 3D-printed devices into experimental workflows.

## Materials and Methods

### Fused filament fabrication (FFF) 3D printing

For 3D print designs we used Autodesk Fusion. For 3D prints we used 2.85 mm filament from Ultimaker and printed it on an Ultimaker S5 printer with the air manager and materials station using Ultimaker Cura slicer software. Although not presented here, we also tested Dremel brand clear PLA and PETG on a Prusa Research printer and did not note any differences in our preliminary toxicity tests. Printer nozzle size was 0.4 mm, wall thickness was set to 1.5 mm. A layer height of 0.15 or 0.2 mm was used with no discernible difference in the parts. The glass print bed was used with glue stick applied to the surface, as an adhesion promoter for all FFF parts. Other print settings in the Cura software were set to default. Unwanted hydration in the filament can lead to imperfections and this is most apparent with clear filament. All filament was kept dry or dried directly before use.

### Low temperature paraformaldehyde sterilization and validation

Items to be sterilized were placed into standard steam/ETO self sealing sterilization pouches (Fisherbrand #0181250), or clean culture petri dishes with covers on but not sealed. A formaldehyde Type 4 multicritical process variable indicator strip (True Indicating CFVG-14) was included inside the pouch or cultureware to monitor the process. We placed the dishes and/or packets into an airtight 3 L container (Snapware) with an open vial of 1 to 4 grams of powdered paraformaldehyde along with an open jar containing about 100 mL of saturated sodium chloride. Water is essential for the process and saturated sodium chloride provides an atmosphere of about 75% relative humidity. The sealed container was placed in an incubator at 37°C for as long as necessary to turn the indicator green. Two days was generally sufficient with the color taking longer to develop in the sealed pouches versus the culture dishes. Once complete we vented the container in a fume hood, the paraformaldehyde vial and jar of saturated sodium chloride are removed and capped until needed again. We did not observe any measurable loss of paraformaldehyde.

If there are deleterious effects on cell viability after the paraformaldehyde sterilization procedure we determined that post processing with ammonia vapor neutralizes most of these negative effects. To post-process with ammonia we place the sterile items, still in their packets/petri dishes, into an airtight container with a beaker containing 2 mL of 3% ammonium hydroxide. The container is then sealed and placed back at 37°C for 24 hours. After the 24 hour incubation the container is aired out in a fume hood and the ammonia treated sterile items are ready for use.

For validation of our low temperature formaldehyde sterilization procedure we used two strains of bacteria, the universally recognized test organism *Geobacillus stearothermophilus* (ATCC strain 7953) and *Escherichia coli* (strain BL21) (19–21). Liquid cultures were shaken in 14 mL polypropylene tubes with *G. stearothermophilus* grown in ATCC Medium 3 at 55°C and *E. coli* grown in LB media at 37°C. For growth on 100 mm bacterial plates the media was made with 1.5% agar. Plates were incubated at 55°C in polyethylene bags commonly used with heat sealers, unsealed but folded over once. 3D print test samples were PLA paddles (8mm x 65mm x 1mm) inoculated with 4 μL of a saturated culture of *G. stearothermophilus* or *E. coli*. The paddles were air dried in a horizontal laminar flow clean bench and placed into standard steam/ethylene oxide self sealing sterilization pouches for use in the sterilization assays. For the 72 hour assay, samples spent 2 hours in the bags prior to incubation with paraformaldehyde, while the 24 and 48 hour samples were in the bags for 8 days prior to incubation with paraformaldehyde. The samples were subjected to sterilization as described above without ammonia vapor neutralization. Control samples, without paraformaldehyde, were placed into a separate sealed container with an open jar of saturated sodium chloride solution and incubated alongside the test samples. Upon completion of the sterilization trial the paddles were aseptically transferred to culture tubes with 2 mL of the appropriate liquid growth medium and incubated with shaking at the appropriate temperature for 48 hours. Any obvious growth in the liquid culture was noted. After the 48 hours of growth in liquid media, 200 μL of culture was spread onto agar plates and incubated for 24 hours at the appropriate temperature. For the 3D print sample to be considered sterile we required the control samples to show growth while the treated samples had no growth. Each strain had 2 biological replicates.

### Cultured cell growth assays with 3D printer filament and adhesives

HL-60 cells were obtained from and grown per the recommendations of ATCC. Briefly, cells were grown in RPMI 1640 medium (Gibco #22400089) with 10% heat inactivated fetal calf serum (Gemini Bio Products #900–108) and an antibiotic antimycotic mixture (Gibco 15240096). Cultures were maintained at 37°C with 5% CO_2_. Cells were kept between 1x10^5^ and 1x10^6^ cells/mL by subculturing every 2 to 3 days. Cells were at 1x10^5^ at the beginning of each assay. For determining cell counts and viability we performed flow cytometry on a BD Accuri C6 Plus flow cytometer. It should be noted that the parent culture can be cleaned up periodically to remove dead cells by density gradient centrifugation with Optiprep (22).

For plastic viability assays we printed plastic disks 12 mm in diameter and 1.5 mm thick with 100% infill. To remove glue stick the disks were cleaned with DI water followed by type 1 water and stored dry until further use. Prior to initiating assays the disks were placed into 24 well tissue culture plates and used neat (non-sterile) or were subjected to paraformaldehyde sterilization with or without ammonia treatment.

For adhesive viability our initial testing used a 14 mm dollop of adhesive on new untreated 18 mm diameter glass coverslips, cured according to the manufacturers recommendations. For UV cure adhesives we illuminated the coverslips from above with a UV lightbox (Fotodyne 3-3035) with 254 nm light at a distance of 20 mm for 5 minutes. After allowing 7 days for full cure the samples were incubated in PBS for 1 hour followed by a 1 hour incubation in complete culture media. The coated coverslips were then placed into a 6 well culture plate with 2 mL of HL-60 cell culture. After approximately 24 hours we removed an aliquot and counted 50 μL of culture by flow cytometry.

Follow up adhesive viability testing was performed by spotting a 10 mm dollop of adhesive, or 8x8 mm square of tape, directly into the well of a 24 well cell culture plate followed by curing for a minimum of 7 days. The adhesive samples were either used neat or processed with paraformaldehyde sterilization and ammonia treatment as indicated. Prerinsing in PBS, when indicated, was done by incubating the samples in PBS at 37°C for 24 hours with one buffer change. 1 mL of HL-60 cell culture was used for each well. Aliquots were removed at 24 and 48 hours to determine the viability by counting 50 μL of culture by flow cytometry. Each adhesive treatment had 3 replicates.

## Results

### Low temperature paraformaldehyde sterilization is suitable for FFF plastics

To enable sterilization of temperature-sensitive 3D-printed components, we developed a simple low-temperature paraformaldehyde vapor protocol performed at 37°C (Fig. 2A schematic). Printed test parts were exposed to paraformaldehyde vapor for defined durations (24–72 hours) in a humidity-controlled container and accompanied by a formaldehyde process indicator strip to provide a visual readout of exposure. We validated sterilization efficacy by intentionally contaminating printed parts with *Escherichia coli* and *Geobacillus stearothermophilus* and assessing recoverable growth following treatment.

**Figure 2:**
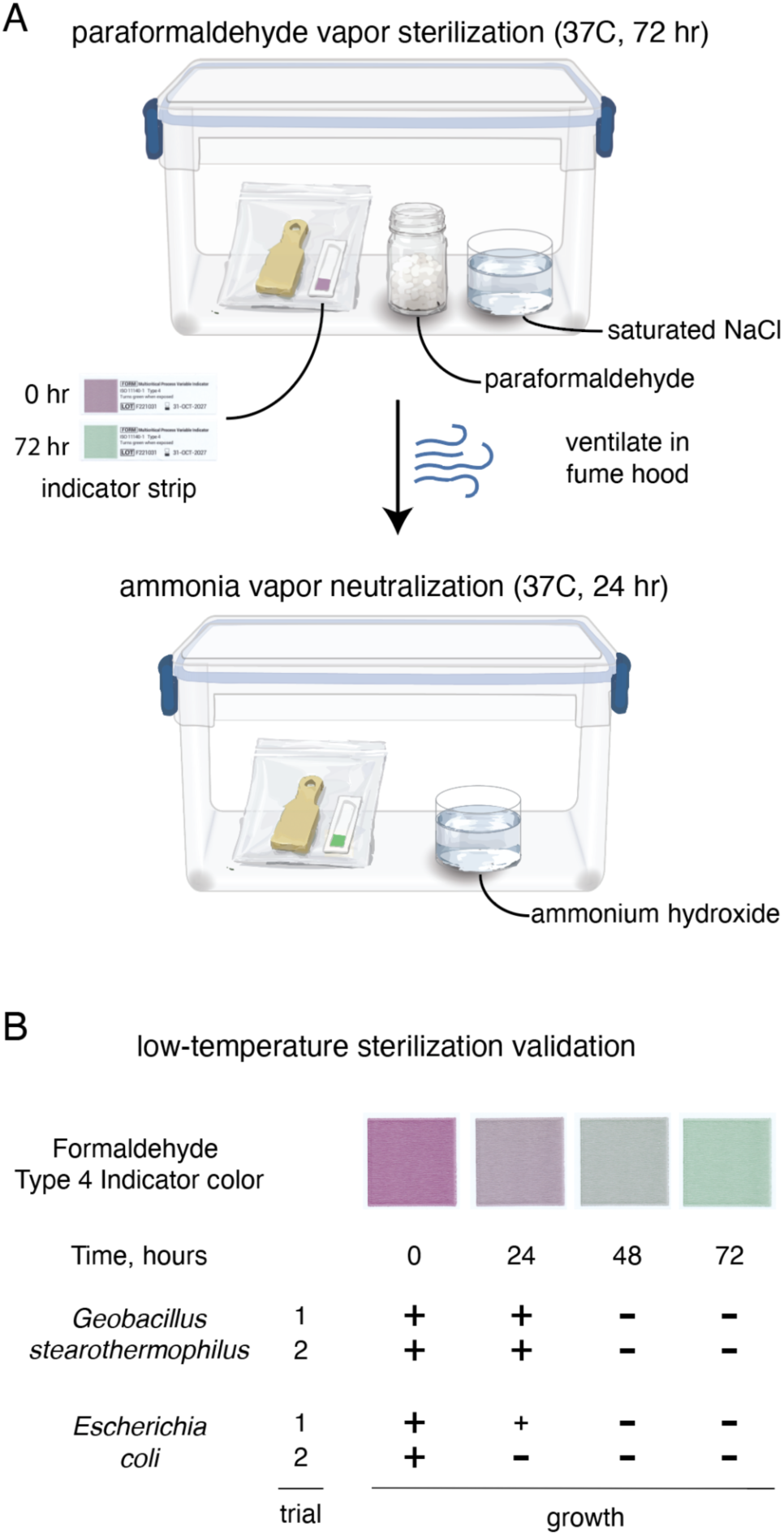
Validation of a low temperature sterilization procedure. (A) Cartoon schematic illustrating the sterilization process utilizing a closed container with paraformaldehyde powder and a humid atmosphere. (B) Validation of sterilization protocol with two bacterial strains dried onto 3D printed PLA plastic sample paddles; extremophile *Geobacillus stearothermophilus* (ATCC 7953) and a standard lab strain of *Escherichia coli*. The assay used two biological replicates with “+” indicating growth and - indicating no growth. Notably, at 24 hours one *E. coli* replicate showed no growth while the other *E. coli* replicate had noticeably less growth indicated by the smaller “+”. The sterilization resistant extremophile was clearly more resistant to sterilization than *E. coli* but is killed before the indicator registers a complete color change.

Using this procedure, *E. coli* and *G. stearothermophilus* contaminated 3D-printed polylactic acid (PLA) plastic paddles showed zero growth after 48 hours of exposure to paraformaldehyde vapor (Fig. 2B). We note that a 24 hour exposure significantly reduced *E. coli* growth, with half of our samples showing no bacterial growth, while *G. stearothermophilus* required the full 48 hour sterilization protocol. In parallel, the indicator strip exhibited a progressive color change with increasing exposure time, shifting from violet (untreated) to gray/purple (24 hour), gray/green (48 hour), and distinctly green by 72 hours, providing a simple visual cue that tracked with effective sterilization conditions.

Sterilization performance required a hydrated atmosphere, and we used a saturated sodium chloride solution which maintains a ∼75% relative humidity while minimizing condensation within the container. We also observed that enclosure and packaging materials influenced exposure of paraformaldehyde vapor. Items placed in closed but unsealed Petri dishes reached the green indicator endpoint more rapidly than items sealed in conventional gas sterilization pouches. Finally, because complex geometries may slow vapor penetration, larger parts or those with deep, narrow cavities may benefit from longer exposure times or from approaches that improve vapor circulation.

### Many plastic filaments used in FFF are safe for short-term biological use

We next assessed growth of human HL-60 cells in the presence of plastics used in routine 3D printing to determine which ones would be best suited for work with cultured cells: polylactic acid (PLA), polyethylene terephthalate (PETG), polypropylene (PP), polycarbonate (PC), thermoplastic polyurethane (TPU), co-polyester (CPE) and nylon. We chose translucent filaments without any colorants to minimize additives and used minimal post-processing, noting that the porous nature of some 3D prints could lead to more additives leaching out over time or contain residual contaminants. To assess potential toxicity, we grew HL-60 cells in the presence of 3D printed disks over 48 hours. Cell growth and abundance was monitored using a flow cytometer (Fig. 3), allowing us to monitor for aberrations in cell morphology, which is reflected in changes in forward and side scatter measurements. Comparisons were made relative to growth of HL-60 cells in the absence of any 3D printed disk.

**Figure 3.**
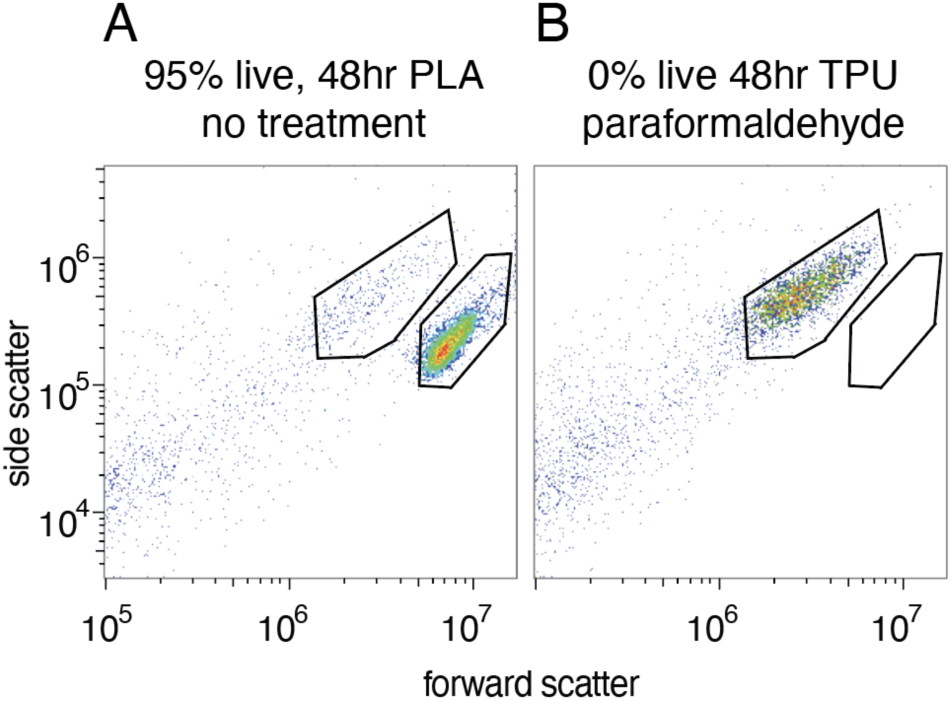
Representative flow cytometry scatter plots and gates used to quantify live and dead cell populations. Forward- and side-scatter profiles are shown for HL-60 cells cultured for 48 h with (A) untreated PLA, yielding 95% live cells, or (B) paraformaldehyde-treated TPU, yielding 0% live cells. The live-cell gate was defined using healthy control cultures whose appearance by routine microscopy was consistent with the population shown in panel A. Scatter-based gating may classify cellular debris as dead cells, potentially overestimating the dead-cell fraction.

We first assessed growth of HL-60 cells with 3D printed disks in the absence of our low temperature sterilization protocol. Encouragingly, most of the plastics did not dramatically alter the relative number of cells quantified after 48 hours of growth (Fig. 4A), suggesting that growth was unaffected by their presence in the cell culture media. This was irrespective of whether we performed a pre-incubation wash of the plastic disks in PBS prior to use. Notably, PLA, PP, and PETG appear safest to use, with no significant difference in cell count relative to control cells that were grown in the absence of a plastic disk. PC, TPU, CPE, and Nylon had small but statistically significant decrease in cell counts relative to the control group. We note that contamination was not observed over this 48-hour period, however, cells were grown in a cell culture media containing a standard antibiotic-antimycotic cocktail (penicillin, streptomycin, and amphotericin B). Sterilization would be required for work with cell culture experiments exceeding 48 hours or where addition of antibiotics and antimycotics is not possible.

**Figure 4:**
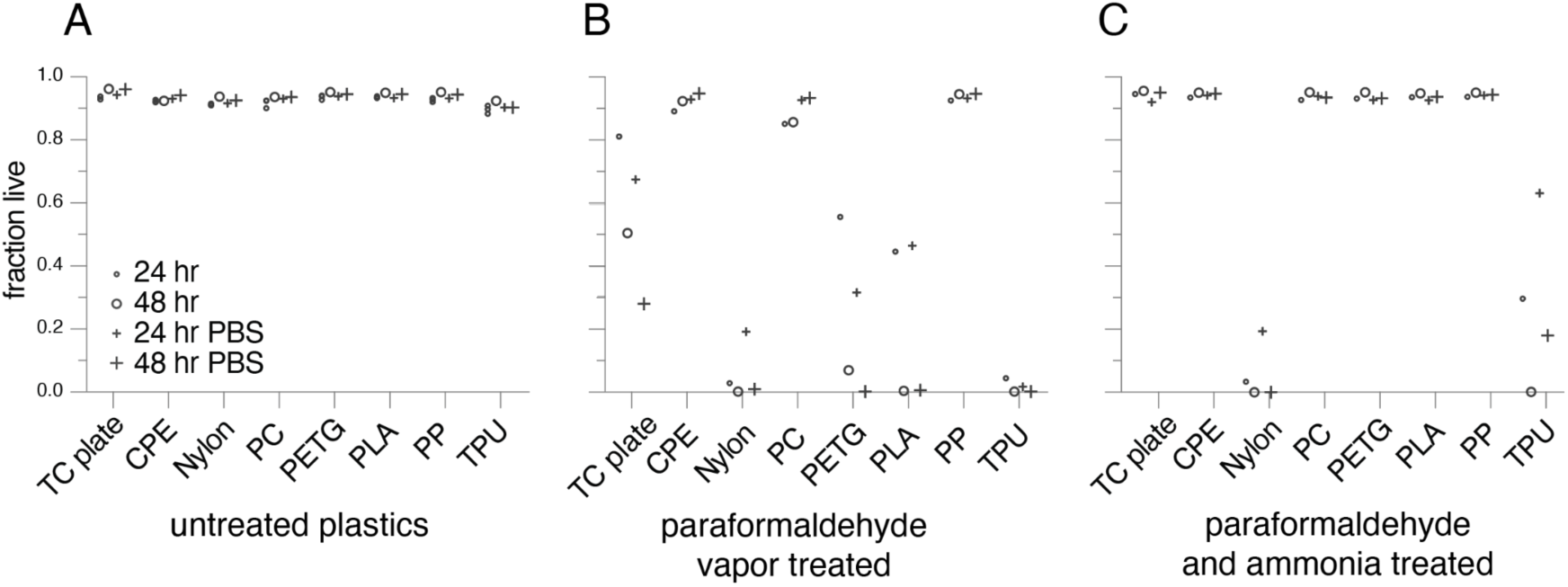
Cytotoxicity of 3D printed plastics without and with parformaldehyde vapor sterilization. HL60 cells, in suspension, were grown with plastic disks utilizing several FFF plastics. The fraction of the surviving cells was determined by flow cytometry after growth for 24 or 48 hours. (A) Cells grew well with most plastics without detectable contamination for up to 48 hours even without sterilization. (B) Low- temperature paraformaldehyde sterilization was deleterious to cell viability with most plastics except for CPE, PC and PP. The increase in cell viability after 24 hours with CPE and PC suggests that any minor residual aldehyde contamination could be overcome. (C) When paraformaldehyde vapor sterilization was followed by ammonia vapor treatment virtually all of the 3D FFF plastics were safe for cell culture. Notable exceptions are nylon and TPU, two polymers that contain amines.

We repeated our cell culture experiments using formaldehyde-treated 3D printed disks to evaluate post-sterilization compatibility. Notably, PP, PC and CPE disks remained nontoxic following formaldehyde treatment (Fig. 4B). However, the other plastics showed a dramatic drop in viable cells, including treatment of our tissue culture (TC) plate without any addition of printed plastic disks. Prior work has shown that formaldehyde-based sterilization can leave residual contamination that may be toxic to cells, and urethanes and nylons in particular have been shown previously to retain high amounts of formaldehyde residue (18). This is consistent with our observations, which included formaldehyde-treated TPU and nylon disks, suggesting that toxic formaldehyde residue remains. To circumvent this, we reasoned that a post-treatment of the plastics with an amine might be sufficient to neutralize any residual formaldehyde. We chose ammonia which has a high vapor pressure and dissipates rapidly. To post-process samples with ammonia vapor, we placed the sterile disks into an airtight container with a beaker containing 3% ammonium hydroxide solution. Plastic disks were incubated at 37°C for 24 hours and then subsequently we reassessed the viability of HL-60 cell growth over a 48-hour period. As shown in Fig. 4C, following ammonia vapor treatment most plastics exhibited no toxicity, with cells exhibiting similar viability as observed with our untreated plastics. Two exceptions to this were the TPU and nylon disks, which continued to exhibit substantial toxicity, with little to no viable cells after 48 hours.

### Many adhesives are toxic to cells when incubated in biological cell culture solutions

Adhesives are important for assembling multipart devices or where a single 3D printed part is not possible. We initially conducted a screen across a selection of adhesives, including several options designed to be safe for medical use or commonly used in microfluidics fabrication. We spotted a patch of adhesive approximately 14 mm in diameter onto 18 mm glass coverslips. The adhesive coated coverslips were placed into individual wells of a 6-well cell culture plate, rinsed in PBS and then cell culture media prior to adding HL-60 cells in fresh media. This approach provided a large, exposed surface area of adhesive and limited post-processing before adding cells, providing a worst-case scenario when using these adhesives. Interestingly many medical grade adhesives exhibited significant toxicity in at least one trial and were not considered further (Table 1). We also observed significant variability between trials when utilizing the UV cure adhesives. We conclude that the designation of “medical grade” does not mean non-toxic with our cell-based assay. In addition, absent commercial-grade UV curing systems, UV cure adhesives are not well suited for fabricated parts designed for cell culture work in the laboratory setting.

**Table 1:** Adhesives ruled out during initial testing. Growth tests were carried out on a 14mm dollop of adhesive spotted onto a glass coverslip followed by curing, aging and rinsing in PBS and culture media. Survival rates were variable and often below 50% relative to a no adhesive control. Of note is the fact that the designation of medical grade (ISO10993 compliant) did not indicate an adhesive was suitable for cell culture.

| Adhesive | Medical | Type |
| --- | --- | --- |
| Loctite 431 | Y | Cyanoacrylate, moisture cure |
| Loctite SI 5031 | Y | Silicone UV cure |
| Loctite SI 5240 | Y | Silicone UV cure |
| Loctite UK M-11FI | Y | Urethane, 2 part |
| Loctite 3106 UV | - | Acrylic, UV cure |
| Krazy Glue All Purpose | - | Cyanoacrylate, moisture cure |
| Loctite 435 | - | Cyanoacrylate, moisture cure |
| Loctite 4311 | - | Cyanoacrylate, UV cure |
| Hardman Double/Bubble | - | Epoxy, 2 part |

We also observed strong chemistry-dependent differences in biocompatibility. Cyanoacrylates and conventional two-part epoxies were uniformly toxic in our assay, with no cell survival after 48 hours, even when marketed for medical use (Table 1). This is consistent with observations in other cell culture systems (23–25). In contrast, silicone-based materials were the most consistently compatible, and we identified both an RTV silicone (Loctite Si 5011 CL) and a silicone adhesive tape (S1001) that were non-toxic in the absence of sterilization and versatile for assembling multi-part devices. Sylgard 184, which is a specific formulation of polydimethylsiloxane (PDMS) designed for use in the electronics industry as a potting compound, is widely used for microfluidics with cultured cells and is safe for use. Lastly, the commonly used Dow Corning high vacuum grease (also sold as DC111 Compound to the food service industry) has been used in cell culture applications for decades and we did not observe toxicity in our tests.

We performed a secondary test designed to better assess compatibility following sterilization. In this format, adhesives and tapes were placed directly into tissue-culture wells, subjected to low-temperature paraformaldehyde vapor sterilization, and then evaluated for effects on HL-60 growth over 24–48 hours. This testing identified additional tapes/adhesives that remained compatible after sterilization and reinforced silicone-based adhesives as the most robust class (Fig. 5). Of note however, among the silicone-based materials, Loctite Si 5011 CL was the only adhesive that was safe without formaldehyde treatment but became toxic after paraformaldehyde exposure. Loctite Si 5011CL, is a low corrosive silicone designed for electronics work that uses a non-corrosive oximino-silane crosslinker in place of the more common acetoxy silane. It is possible that this amine containing crosslinker is responsible for the adhesive’s reactivity with formaldehyde. Indeed, subsequent ammonia vapor neutralization followed by a PBS rinse completely restored compatibility with cells (Fig. 6). Together, these data indicate that while silicone-based adhesives are generally robust to paraformaldehyde sterilization, specific formulations can become cytotoxic and may require post-treatment to restore compatibility.

**Figure 5:**
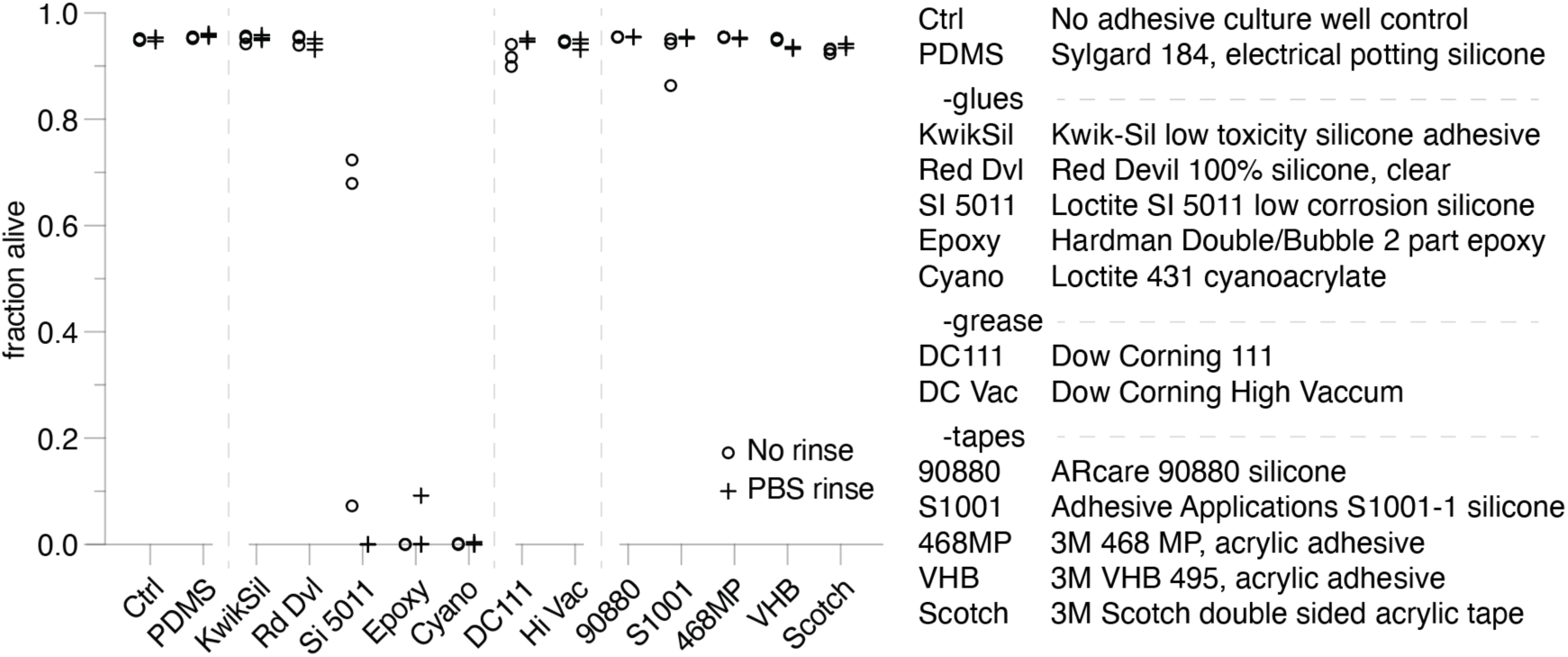
Cytotoxicity of adhesives to cell culture with parformaldehyde vapor sterilization. Adhesive viability tests utilizing a 10 mm dollop of adhesive in the well of a 24 well culture plate subjected to paraformaldehyde sterilization without ammonia neutralization indicates many adhesives are generally safe for work with cultured cells. Of note is the cyanoacrylate was a medical grade product and the Red Devil silicone is a consumer product. Each experiment has a total of 3 replicates, some with overlapping data points.

**Figure 6:**
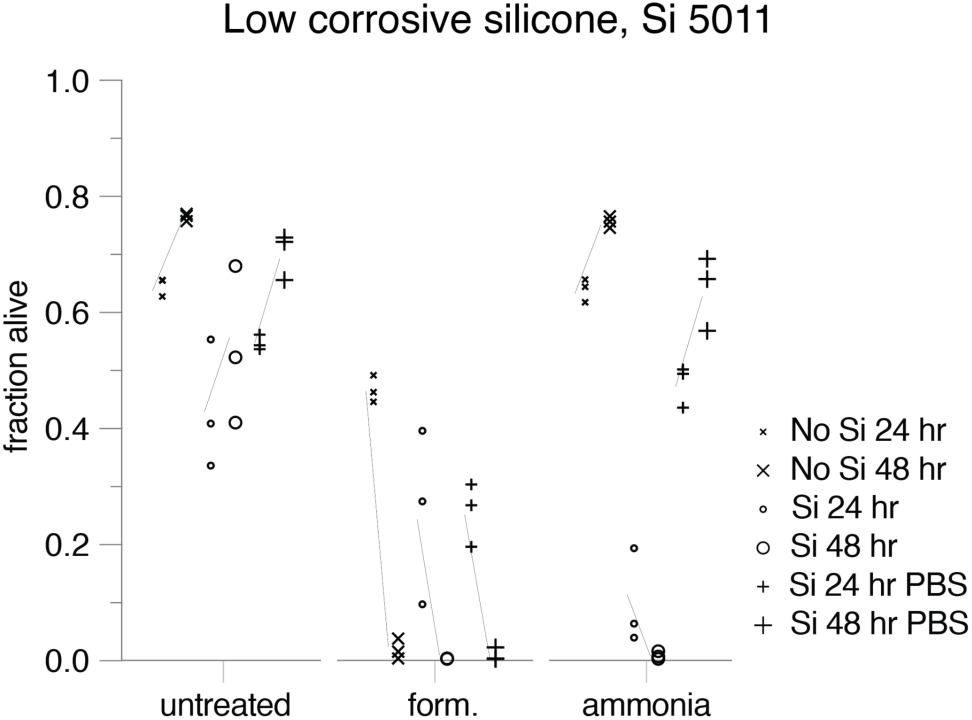
Cytotoxicity of Loctite Si 5011 low corrosive silicone without and with parformaldehyde vapor sterilization. Trend lines indicate that cells can proliferate in the presence of the adhesive without sterilization or with sufficient neutralization and rinsing post sterilization. This silicone adhesive is low corrosive for metals, replacing a common acetoxy silane curing agent with an oximino silane. While ammonia treatment restored compatibility with cells following parformaldehyde sterilization, we note that the fraction alive for the no-silicone adhesive control is lower than in the previous figure, due to the higher cell culture passage number.

## Discussion

FFF-based 3D printing is a versatile, iterative, and inexpensive fabrication approach that can be applied to many different experimental needs. We have developed a low-temperature sterilization strategy and validated its compatibility for biological cell culture across an array of FFF plastics and adhesives. This work also identifies important considerations in the choice of fabrication materials where sterility and biocompatibility are required.

In most biology laboratories, sterilization of media and equipment is performed using autoclaves that maintain temperatures and pressures incompatible with many FFF plastics. While cured photopolymers used in stereolithography-based 3D printers can generate printed parts that tolerate high temperatures, they often exhibit poor biocompatibility, likely through leaching of toxic compounds into culture media (13, 26–28). Similarly, we suspect that the toxicity observed with some UV-cured adhesives may result from incomplete curing and leaching of residual chemicals into the cell culture media. Other sterilization approaches, including ionizing radiation, ethylene oxide, and low-temperature steam (occurring at 70°C) and formaldehyde, require equipment that is not readily available in many research laboratories (26, 27). We therefore explored low-temperature gaseous sterilization methods (28). The protocol described here is a modification of low-temperature steam formaldehyde sterilization procedures previously developed for medical applications (18). We validated our method utilizing the test organism for low-temperature steam and formaldehyde sterilization processes specified in ISO 11138-5. We believe that our method complies with much of the requirements for sterilization of health care products in ISO 14937 with the major exception that we only tested against bacteria.

Sterilization at 37°C ensures compatibility with most FFF assemblies. We also evaluated gaseous chlorine dioxide sterilization, which successfully sterilized *E. coli*- contaminated printed plastic but produced a distinct green discoloration of the material (data not shown), and therefore was not pursued further.

Among the plastics tested, PLA was the easiest filament to print reliably and rarely required additional printer maintenance. PETG prints appeared more watertight under pressure using our current workflow. We note that PLA components do not withstand prolonged exposure (weeks) to aqueous environments and should generally be considered disposable.

Silicone RTV adhesives, including the consumer grade one we tested, and silicone-based tapes were the most reliable assembly materials across both untreated and post-sterilization conditions and may therefore serve as useful starting points for fabrication of multi-part devices. The acrylic tapes we tested were also very good with the viability of the 3M 468 MP being on par with the silicone adhesive based tapes. This is significant given the potential difficulty in obtaining compatible double sided silicone tapes. It’s worth noting that our adhesive tests exposed a large surface area of adhesive to the cell culture media. For situations where adhesives have minimal exposure to the cell culture media, some “incompatible” adhesives may still be usable.

When sterilization is required, materials with amines such as TPU, nylon and adhesives with amine crosslinkers should be avoided because of their propensity to retain aldehyde-associated toxicity. For several other materials, ammonia vapor neutralization was sufficient to restore compatibility following paraformaldehyde vapor treatment. One variable we did not test was the effect of outgassing time, post formaldehyde treatment, on toxicity. It’s possible that negative effects of paraformaldehyde vapor sterilization will be mitigated by prolonged post treatment outgassing prior to use without ammonia neutralization.

While we limited our assessment of biocompatibility to short-term viability and growth assays using HL-60 cells, this approach was sufficient to identify a range of plastics and adhesives suitable for cell culture applications. For particularly sensitive cell models or long-term culture experiments, additional validation using complementary assays that directly assess proliferation, apoptosis, or other cellular responses may be warranted (29, 30). On the other hand, we identified several adhesives that are safe for medical use and ISO 10993 compliant but were toxic in our viability assay. These results reflect the specific requirements of cell culture applications rather than general medical biocompatibility.

## Conclusions

We developed a low-temperature paraformaldehyde vapor sterilization protocol compatible with common FFF plastics and validated its efficacy using bacterial challenge assays. Effective sterilization was achieved without exposing printed components to temperatures that would deform many commonly used materials. Among the plastics tested, PLA, PETG, and polypropylene exhibited strong compatibility with cultured HL-60 cells, while silicone-based adhesives and tapes along with acrylic tapes were the most consistently biocompatible assembly materials. For several materials, residual toxicity associated with paraformaldehyde treatment could be mitigated through ammonia vapor neutralization. Together, these findings provide practical guidance for fabrication, sterilization, and deployment of custom 3D-printed hardware in cell biological research.

## Future Perspective

As 3D printing becomes increasingly integrated into biological research workflows, low-cost fabrication methods are likely to accelerate the development of custom microscopy chambers, cell migration devices, microfluidic systems, and laboratory automation hardware. Future work could expand compatibility testing to additional filament formulations, primary cell models, and longer-term culture experiments. The low-temperature sterilization strategy described here is easy to implement and could be useful where more traditional methods are not achievable such as remote research facilities. It may also prove useful for sterilizing conventional laboratory plastics, cell culture plastics and mixed-material assemblies that cannot tolerate autoclaving, extending its utility beyond 3D-printed components.

## Article Highlights

- A simple low-temperature paraformaldehyde vapor sterilization protocol was developed for fused filament fabrication (FFF) components and validated using bacterial challenge assays.
- Effective sterilization of 3D-printed parts was achieved under conditions compatible with common FFF plastics.
- PLA, PETG, and polypropylene exhibited good compatibility with HL-60 cell culture, while TPU and nylon showed reduced compatibility following sterilization.
- Residual toxicity associated with paraformaldehyde sterilization could be mitigated for many materials by post-treatment with ammonia vapor.
- Silicone-based adhesives and tapes were the most consistently compatible materials for fabrication of cell culture devices.
- The combination of low-temperature sterilization and compatible fabrication materials provides a practical framework for rapid prototyping of custom hardware for cell biological research.

## Authors’ Contributions

Matthew J. Footer: Conceptualization, Methodology, Investigation, Writing: original draft, Writing: review & editing.

Nathan M. Belliveau: Conceptualization, Investigation, Writing: original draft, Writing: review & editing.

## Acknowledgements

We thank Thomas Riha at True Indicating for helpful discussions on formaldehyde sterilization. We also thank Maya Ozo for helpful literature recommendations and Mitch Sanders for editorial comments. M.F. is supported through research funding from the Howard Hughes Medical Institute. N.M.B is supported by a grant from the National Institutes of Health (R00GM147355). ChatGPT was used to make part of the sterilization process illustration in Fig. 2A.

This article is subject to HHMI’s Immediate Access to Research Policy. A preprint version is freely available under a CC BY 4.0 license.

